# Running to remember: The effects of exercise on perineuronal nets, microglia, and hippocampal angiogenesis in female and male mice

**DOI:** 10.1101/2023.11.24.568577

**Authors:** Madeleine G. Maheu, Noah James, Zach Clark, Alex Yang, Ridhi Patel, Shawn M. Beaudette, Rebecca E.K. MacPherson, Paula Duarte-Guterman

## Abstract

Exercise is accepted as a positive health behaviour; however, the mechanisms of exercise on neuroprotection and cognitive health are not completely understood. The purpose of this study was to explore the neurobiological benefits of chronic treadmill exercise in female and male mice through its role in microglial content and morphology, cerebral vascularization, and perineuronal net (PNN) expression. We further examined how these neurobiological changes relate to spatial memory outcomes. Adult mice were assigned to a sedentary or treadmill exercise group for eight weeks. During the final week, all mice were trained on a spatial memory task (Barnes maze) and brains were collected for immunohistochemistry. Exercised mice made fewer errors than sedentary mice during the first two days of training and probe trial. Females, regardless of exercise training, made fewer errors during Barnes maze training and demonstrated a greater frequency of spatial strategy use compared to males. Exercised mice, regardless of sex, had fewer PNNs in the dentate gyrus of the hippocampus compared to sedentary controls. The number of PNNs in the dorsal dentate gyrus was positively correlated with total errors during training. During the probe, greater errors correlated with more PNNs among the exercised group only. Microglia count and cerebral vascularization were not affected by exercise, although proportions of microglia type (ameboid, stout/thick, and thick/thin) were regulated by exercise in the ventral dentate gyrus. We conclude that exercise decreases PNNs in the dentate gyrus in both sexes and this may be related to better spatial learning and memory.

## Introduction

Exercise is widely accepted as beneficial for health, positively affecting almost every tissue in the body including the brain (Pedersen & Saltin, 2015). However, most literature pertaining to the neurobiological benefits of exercise has been derived from studies utilizing male animal models and sex differences have often been overlooked in studies examining the benefits of exercise on brain health. The mechanisms of how exercise benefits the brain and cognitive health in both sexes are less well understood (Barha et al., 2019). Possible neuroprotective mechanisms of exercise include regulation of inflammation and plasticity through microglia and perineuronal net (PNN) expression, and regulation of angiogenesis. The purpose of this study was to examine the propensity for exercise to influence spatial learning and memory and cellular mechanisms in the brain via neuroplastic, neuroimmune, and neurovascular changes in adult female and male mice.

Perineuronal nets (PNNs) are collections of extracellular matrix molecules, which have a primary role in synaptic plasticity through the stabilization of synapses between neuronal somas and proximal dendrites (Celio et al., 1998). PNNs have a role in neuroprotection and memory formation through the regulation of neuronal excitatory and inhibitory balance, and the removal of PNNs results in decreased synaptic inhibition (Lensjo et al., 2017; Khoo et al., 2019) and improvement in recognition memory (male mice; Gogolla et al., 2009; Romberg et al., 2013). Additionally, PNNs repel neurites and growth cones (Xiao et al., 1996), thereby preventing new synaptic connections (Celio et al., 1998). Synapses with fewer PNNs are thought to be less stable and more plastic than those with greater PNNs (Bozzelli et al., 2018). Although there has been little research on the effects of exercise on PNN distribution, previous studies have found that the number of PNNs is reduced in response to volunteer wheel running. Smith and colleagues (2015) demonstrated lower PNN density in the CA1 region of the hippocampus of adult female rats in response to six weeks of voluntary wheel running.

Additionally, voluntary wheel running (for 4 weeks) also reduced PNN numbers in the dentate gyrus in male C57BL/6J mice (Briones et al., 2021). A decrease in PNN expression with exercise may indicate enhanced plasticity, which could influence hippocampal-dependent memory formation, however memory has not been assessed in previous studies. In addition, the relationship between PNNs and memory may be complex, as it has also been postulated that PNN formation is essential for the consolidation of memories (Shi et al., 2019). Moreover, digestion of PNNs (through chondroitinase ABC treatment) increased susceptibility to fear memory erasure (Gogolla et al., 2009). Collectively, changes in PNNs can result in a variety of learning outcomes suggesting a complex relationship between the precise balance of PNNs and their involvement in memory.

Microglia, the immune cells of the central nervous system, have been implicated in learning and memory (Czerniawski et al., 2014; Wadhwa et al., 2017; Nguyen et al., 2020) such as maintenance of synaptic connections and synaptic pruning (Paolicelli et al, 2011), neuronal survival and activity (Sierra et al., 2010), axonal trogocytosis (Lim & Ruthazer, 2021), phagocytosis of cellular waste (Ito et al., 2007), and secretion of immune factors (Zhang et al., 2011; Zhou et al., 2012; Jana et al., 2014, Chen et al., 2019; Koss et al., 2019). In both human and mouse models of neurological disease, PNN components have been found in phagocytosed material within microglia (Crapser, 2020a). Brain regions with high PNN intensities tend to have fewer microglia (Barahona et al., 2022) and decreases in microglia through pharmacological intervention has been associated with increased PNNs in both neurological disease models (Crapser et al., 2020ab) and in healthy adult mouse brains (Liu et al., 2021). Recent research has also directly implicated microglial secretions of inflammatory proteins in reducing PNN integrity (Wegrzyn et al., 2021). Taken together, these studies provide support for the concept that microglial activation is involved in PNN disassembly. However, to date, there have been no studies investigating the relationship between microglia and PNN regulation with respect to exercise intervention. In mouse models of neurological disease (Alzheimer’s disease, experimental autoimmune encephalomyelitis, and Parkinson’s disease) there is evidence that exercise interventions decrease microglia number in the substantia nigra pars compacta, striatum, and lumbar spinal cord and produce a hiatus of ameboid activity through a reduction in CD11b expression in males (Sung et al., 2012) and females (Benson et al., 2015). Aerobic exercise-induced neural precursor cell activity is mediated by microglia, and the removal of microglia can eliminate the neurogenic gains from exercise in female mice (Vukovic et al., 2012). Altogether this would suggest that the benefits of exercise on cognitive health are in part produced by the optimization of microglia function; however, it is not clear whether exercise decreases inflammation in healthy wildtype female and male mice. Previous literature also suggests that microglia may regulate PNN expression through inflammatory breakdown of the extracellular matrix (Venturino et al., 2021), and exercise may upregulate the release of inflammatory factors responsible for PNN degradation (Nishijima et al., 2015). Microglia and PNN function can interact, therefore in this study we examined both endpoints following an exercise regimen in females and males.

Exercise is well-known for increasing blood flow through angiogenesis in skeletal muscle (Prior et al., 2004) but less is known about changes that occur in the brain (Pereira et al., 2007; Clark et al., 2009; Mortland et al., 2017). Angiogenesis occurs in response to signaling mediators or increases in tissue size and energy expenditure (activity). Tissues that increase in size or energy expenditure require enhanced exchange with vasculature for oxygen, nutrients, waste processing, and signaling molecules (Prior et al., 2004). In the brain, voluntary wheel running increases blood volume in the mouse dentate gyrus in the hippocampus (sex not reported), and these changes have been associated with increased adult neurogenesis (Pereira et al., 2007). Additional work has demonstrated that exercise enhances blood flow through angiogenesis in the rodent brain by concomitantly acting on multiple stimulating factors (Pereira et al., 2007; Creer et al., 2010; Morland et al., 2017). For example, chronic high intensity interval exercise increases capillary density in the hippocampus in mice, particularly in the dentate gyrus (mixed sexes in combined analysis; Morland et al., 2017). Areas such as the hippocampus, specifically the dentate gyrus (Pereira et al., 2007; Clark et al., 2009), undergo the greatest vascular changes compared to the cerebral cortex where no changes have been observed (Morland et al., 2017). Furthermore, voluntary wheel running increases vascular density in the dentate gyrus and improves performance on the Morris water maze in male mice (Clark et al., 2009). However, these results are strain dependent, observed in C57BL6J but not B6D2F1/J mice despite neurogenic gains (Clark et al., 2009). Overall, there is evidence that brain angiogenic processes increase in response to exercise (Pereira et al., 2007; Clark et al., 2009; Morland et al., 2017), however this may depend on strain while sex differences have not been examined.

Despite the abundance of research regarding the effect of exercise on various neurophysiological measures, there have been fewer studies focusing on exercise-related sex differences in relation to spatial memory and cellular mechanisms. With this gap in knowledge, the purpose of this study was twofold: (1) to characterize the effects of exercise on spatial memory and dentate gyrus levels of, PNNs, microglia, and angiogenesis; and (2) to determine whether there are any sex differences with respect to these measures in healthy adult mice.

## Materials and Methods

### Experimental Design

This experiment was approved by the Research Ethics Board at Brock University (#21-0-03) and followed the guidelines of the Canadian Council on Animal Care. Young-adult (10-weeks-old) C57BL/6J mice (24 males and 24 females) were obtained from Jackson Laboratory. Mice were housed in groups of four, by same sex and condition, kept on a 12/12h light/dark cycle (lights on at 08:00), and had ad-libitum access to water and food (standard mouse chow, 2014 Teklad global, 14% protein rodent maintenance diet, Harlan Tekland, Mississauga, ON). Mouse body mass and food intake were tracked on a weekly basis. Weekly food intake was measured by cage and an average was calculated per mouse. Mice were housed at a temperature of 22-24°C for the duration of the study.

### DXA scan

A small animal dual-energy X-ray absorptiometry (DXA) scanner (OsteoSys InSIGHT, Scintica) was used to non-invasively measure body composition in anesthetized mice (vaporized isoflurane 5% in O_2_). To ensure minimal anaesthetic exposure, animals remained asleep for only the length of the scanning process, which was less than one minute per animal. DXA scans were conducted to elucidate possible brain-body changes, such as determining if body composition changes occurred alongside neurobiological changes. Outcomes included lean and fat mass and bone mineral content and density. Percent lean and fat mass were calculated using total body mass. DXA scans were taken at the beginning (baseline) and the end of the 8-week intervention.

### Exercise by Treadmill Running

One week after arrival to the animal facility, all mice were exposed to the treadmill for two 5-minute habituation periods in which the mice explored the stationary treadmill, and two 10-minute acclimatization periods in which the mice ran on the treadmill at a speed of 15 m per minute and an incline of 5%. Habituation and acclimatization occurred during the light cycle between 9 and 11 am and took a total of two days, with each day consisting of one habituation period followed immediately by one acclimatization period. Mice were assigned to either the exercise (n=12/sex) or sedentary control (n=12/group) condition.

The training protocol began 72 hours after the last acclimation session. Mice were exercised on two rodent-sized six-lane treadmills for one hour per day, five days per week, for 8 weeks. The training was progressive to consistently achieve moderate exertion of ∼65% VO_2_ max (Fernando et al., 1993; McFarlan et al., 2012), beginning at 20 m/min at 15% incline and ending at 25 m/min at 25% incline for the final two weeks. Exercise sessions occurred between 9:00 am and 11:00 am. One of the female mice was taken out of the exercise group 10 days into the study due to an injury while running was not included in any analyses. During all exercise sessions, the mice in the sedentary group were taken into the treadmill room. This intervention was chosen as it has been widely used to induce changes in mitochondrial content in skeletal muscle and adipose tissue, serving as a reliable indicator of exercise-induced physiological adaptations thereby indicating that the body has experienced effects that are consistent with exercise training (Castellani et al. 2014, Snook et al., 2016). Our group has further demonstrated that this exact protocol results in higher brain BDNF content and improved recognition memory in young male mice and that this occurred in the absence of differences in body mass and food intake (Baranowski et al., 2023).

### Barnes Maze Testing

During the last week of the exercise intervention, both the exercise and sedentary control mice were trained on a spatial memory task using the Barnes maze (height 44 cm, diameter 122 cm, 20 holes). This test involved three phases, a habituation phase, a training phase, and probe phase, all of which were adapted from Barnes (1979). Visual cues were placed around the room and remained in the exact same location and orientation for the duration of testing. During the habituation phase to the Barnes maze, each mouse was individually placed in a clear upside-down beaker in the centre of the maze and guided around the inner perimeter for 30 seconds, ending at the escape hole. They were then allowed to explore the escape hole for 3 minutes within the confines of the beaker. At the end of the three minutes, they were nudged through the escape hole into the escape box (13.5 x 7.75 x 5.75 cm) and allowed to explore the box for one minute. The mouse was then removed from the room and returned into their home cage. Barnes maze training began 24 hours after habituation. Mice were placed in the centre of the maze under a beaker. The beaker was lifted, and mice were allowed to explore for two minutes. If they found the hole within those two minutes, they were left to explore the escape hole for one minute. If they did not find the escape hole, they were guided to it and put inside the escape box to explore for one minute. After exploring the escape box, the mice were removed from the room and returned to their home cage. Training took place every day (one trial per day) over five days. Mice completed this task in a different order every day of training. The probe took place 48 hours after the final training session. Mice were placed in the centre and given three minutes to explore the maze without the escape box present. At the end of the three minutes, the mouse was removed from the room and returned to their home cage. Testing took place at the same time every day (between 1-4 pm), 3 hours after the 1-hour treadmill running and was consistent for all mice. All sessions (training and probe) were recorded via video. All materials were disinfected between animals using peroxiguard (1:40 dilution).

### Behavioural Analysis

Overhead videos were taken (Sample Rate: 30Hz; Resolution 1280×720 pixels), and animal tracking was performed using DeepLabCut (DLC v2.2). Specifically, a custom DLC model was trained using user inputs to resolve the locations of all Barnes Maze Targets and the Nose and Tail Base of tracked mice. The DLC model (ResNet 50 architecture) was trained using 20 random frames from 10 input videos for a total of 500,000 epochs, resulting in a mean testing error of 2.35 pixels across all tracked positional landmarks. Tracked XY positional data from DLC were further analysed using custom MATLAB (v2022a) script to scale to real-world units, and to generate discrete performance parameters. Specifically, latency to find the target hole, number of errors (incorrect target visits), number of repeat errors (incorrect repeat visits to target holes), cumulative path length, and average speed, were measured for the training sessions and calculated using MATLAB. An error was defined as approaching an incorrect target hole within a 5 cm radius from the hole centre-point, as measured by the animal tracking software. Time spent in each quadrant and within the vicinity of the target (defined as 5 cm from target hole) were added to the probe analysis. Search strategy was categorised manually while blinded to treatment and sex using an output image of the path taken by each mouse. The following search strategies were included in the analysis: random, serial, random/serial, and spatial. A random strategy was categorised as unsystematic and localized searches of holes by crossing through the centre of the maze (Rosenfield & Ferguson, 2014). A serial strategy was categorised as a systematic search of at least four consecutive holes (Rosenfield & Ferguson, 2014). A random/serial strategy was categorised as having elements of both random and serial strategies (O’Leary et al., 2011). Finally, a spatial strategy was categorised as direct navigation to the correct quadrant, with the exception of crossing the maze centre up to one time, with 3 or fewer errors (Rosenfield & Ferguson, 2014).

### Tissue Collection and Processing

Immediately after the probe, endpoint DXA scans were performed. The day following the probe and 72 hours after the last exercise bout, mice were deeply and continuously anaesthetized with isoflurane to the extent of reflex loss. Brains were perfused with ice-cold 0.9% saline followed by 4% paraformaldehyde in PBS (phosphate buffer saline). Brains were extracted, stored and post-fixed in 4% paraformaldehyde in PBS for 24 hours at 4 °C, after which they were cryoprotected through storage in 30% sucrose in PB solution at 4 °C. Adrenals were collected and weighed as an indicator of stress. Using a freezing sliding microtome (SM2010R, Leica), brains were coronally sectioned in 40 µm increments across the extent of the hippocampus (collected in seven series). Sections were stored in an antifreeze solution containing 0.1 M PBS, 30% ethylene glycol, and 20% glycerol at -20°C.

### Immunohistochemistry

Iba-1 and Wisteria floribunda agglutinin (WFA) were used to visualize microglia and perineuronal nets, respectively and both stains were used on the same sections. Free-floating sections were rinsed three times for 10 min in 0.1 M phosphate buffer saline (PBS; pH = 7.4) in between each step listed below unless otherwise stated. Sections were blocked with 3% normal goat serum (NGS) and 0.3% Triton-X in PBS for 45 minutes. Sections were incubated with rabbit-anti-Iba-1 primary antibody (1:1000, Wako Chemicals; Cat. No) and WFA (1:400; Sigma; Cat. No) for 23 hours at 4 °C. Sections were rinsed in PBS and then incubated with secondary antibodies goat anti-rabbit Alexa 488 (1:200; ThermoFisher) and streptavidin Alexa 568 (1:400; ThermoFisher) for 20 hours at 4 °C. Sections were rinsed with PBS, counterstained with DAPI, mounted onto frosted microscope slides, and cover slipped with using polyvinyl alcohol with DABCO (PVA-DABCO; Sigma-Aldrich).

On another collection of sections, CD31 was used to visualize the vasculature following the protocol by Rust and colleagues (2020). Sections were washed with 0.1M PB and then incubated with blocking and permeabilization solution (TNB, 0.1% TBST, 3% normal goat serum) for 30 minutes at room temperature shaking. Sections were rinsed with phosphate buffer three times for 10 minutes each and incubated with primary antibody (rat anti-mouse CD31, BD Biosciences, 550274, 1:100) overnight at 4°C. The next day sections were rinsed once to remove excess antibody and then three times with phosphate buffer for 10 minutes each. Sections were then immediately incubated with secondary antibody (goat anti-rat Alexa 488, 1:200, ThermoFisher) for 2 hours at room temperature, and then overnight at 4°C. All antibodies were diluted in the blocking and permeabilization solution (TNB, 0.1% TBST, 3% normal goat serum). Sections were washed, counterstained with DAPI, mounted onto microscope slides, and cover slipped using PVA-DABCO.

### Microscopy Analysis

A researcher blinded to experimental conditions imaged and analysed Iba-1+ cells, WFA and CD31 expression. Sections were imaged using an epifluorescence microscope (Olympus BX53) at 200X magnification using the same exposure settings for all samples. The dorsal and ventral hippocampi were analyzed separately to account for the differential functions of these regions, thereby underlying different behaviours (Kheirbek & Hen, 2011). For each brain, images were taken of four sections of the dorsal and three sections of the ventral hippocampus. Ventral was designated as any section that appeared posterior to Bregma -2.92 (Paxinos & Franklin, 2001). ImageJ Fiji was used for image analysis (Schindelin et al., 2012). A region of interest (0.4597 mm x 0.3448 mm, 0.1585 mm^2^) was created around the granule cell layer and hilus regions of the dentate gyrus (Figure 4A), from which WFA, Iba-1 and CD31 expression were counted. We always applied the region of interest in the area where the suprapyramidal blade meets the infrapyramidal blade of the granule cell layer. This was chosen so that we could be consistent between images and individuals. The same region of interest (shape and size) was applied to all images and samples. For each marker, counts were average from four dorsal and three ventral sections per individual. In addition, Iba-1 morphology was categorized into three groups: ameboid, stout/thick, and thick/thin, similar to previous work (Duarte-Guterman et al., 2023; Eid et al., 2019). Categorization was based on the branching of the processes, with an ameboid morphology having no processes, a stout/thick morphology having one to two processes close to the cell body, and a thick/thin morphology having several thin processes extending outwards from the cell body (Figure 5A-C). CD31-labelled vessels were also analysed by counting number of branches per vessel and by length of vessel. Vessel length was measured as the longest traceable entity for each vessel within the region of interest.

### Statistical Analyses

GraphPad Prism (version 8.4.3.686) was used to run all statistical tests. ANOVA assumptions of normality and homogeneity of the variance were fulfilled by Shapiro-Wilk and Brown-Forsythe tests (p > 0.05) as well as visual inspection of QQ and homoscedasticity plots, respectively. For repeated designs, the Greenhouse-Geisser correction was used if the assumption of sphericity was not met. ANOVAs using sex (female, male) and treatment (sedentary, exercised) as between subject factors were performed on the adrenal to body mass ratio, total food intake, total errors, total repeat errors, latency, distance traveled, speed, time spent in the correct quadrant, and time spent near the escape hole during training and the probe session. Two-way independent ANOVAs were also performed on post morphometric parameters and change values in morphometric parameters. Repeated measures ANOVAs with days/weeks as the within subjects’ factor and sex and treatment as between subject factors were performed on pre-to-post body mass, latencies, number of errors, and number of repeated errors across training days. For the neurobiological measures, ANOVAs were performed on Iba-1, WFA, and CD31 expression, using sex and treatment as between subject factors. The dorsal and ventral dentate gyri were analysed separately. When a main effect or interaction was found to be significant, a post-hoc Bonferroni comparison was used for all post comparisons. Pearson *r* correlations were performed on WFA counts with behavioural measures of errors and latencies during Barnes maze training and the probe session. Sedentary and exercise groups were analyzed separately in correlation analyses to account for any potential modulating effects of exercise on PNN expression in relation to memory outcomes. We did not correct for multiple correlations as adjusting the alpha value can be overly conservative and increases the chances of Type II errors. These correlations are exploratory and present potential mechanisms to pursue in the future. We report significance differences (*p* ≤ 0.05) and trends (p ≤ 0.08).

## Results

### Exercise did not alter body composition

All changes in morphometric parameters with sex and exercise are summarized in Table 1. Exercise did not significantly affect body composition although there were trends - specifically the reduced change in fat mass among the exercised group compared to the sedentary controls (main effect of exercise: *F* (1, 43) = 3.625, p = 0.0636)) and the increase in adrenal mass with exercise in females (interaction between exercise and sex: (*F* (1, 33) = 3.383, *p* = 0.0749)). Regardless of exercise, we found significant sex differences in body and bone composition across nearly all morphometric parameters and in caloric intake. Food intake was higher in males compared to females (p<0.0001) but there was no effect of exercise on this parameter.

**Table 1.** Effects of exercise training and sex on morphometric parameters, adrenal mass, and food intake in male and female mice.

|  | Male |  | Female |  | ANOVA, p |  |  |
| --- | --- | --- | --- | --- | --- | --- | --- |
|  | Sed. | Ex. | Sed. | Ex. | Ex. | Sex | Intx |
| Body mass (g) | 32.03±2.03 | 30.95±2.22 | 22.32±1.28 | 22.45±0.9 | >0.3475 | **<0.0001** | >0.2295 |
| Fat mass (g) | 3.23±0.89 | 2.83±0.50 | 1.80±0.22 | 1.79±0.22 | >0.2084 | **<0.0001** | >0.2145 |
| % fat mass | 10.1±2.4 | 9.1±1.4 | 8.1±0.9 | 8.0±1.1 | >0.3087 | **0.0017** | >0.3488 |
| Change in fat mass | 0.79±0.57 | 0.54±0.54 | 0.34±0.32 | 0.36±0.14 | >0.1044 | **0.029** | >0.8482 |
| Change in fat % | 0.010±0.015 | 0.002±0.017 | 0.004±0.020 | 0.006±0.014 | **0.0636** | >0.2698 | >0.8082 |
| Lean mass (g) | 28.30±1.82 | 27.62±1.99 | 20.13±1.16 | 20.26±0.96 | >0.5493 | **<0.0001** | >0.3373 |
| % lean mass | 88.4±2.5 | 89.3±1.4 | 90.2±1.2 | 90.2±0.9 | >0.3435 | **0.0057** | >0.3654 |
| Change in lean mass (g) | 4.21±1.40 | 4.62±0.79 | 3.12±0.83 | 2.56±1.14 | >0.7975 | **>0.0001** | 0.1274 |
| Change in lean % | 0.011±0.015 | 0.003±0.018 | 0.003±0.015 | 0.004±0.021 | >0.1458 | 0.1565 | 0.8830 |
| *Adrenal/body mass ratio | 1.175±0.301 | 1.02±0.353 | 1.328±0.393 | 1.66±0.515 | >0.5100 | **0.0054** | **0.0749** |
| Bone mineral density | 0.072±0.002 | 0.074±0.002 | 0.070±0.003 | 0.070±0.002 | >0.5267 | **<0.0001** | >0.2026 |
| Bone mineral content | 0.502±0.037 | 0.494±0.042 | 0.392±0.027 | 0.397±0.018 | >0.9491 | **<0.0001** | >0.4907 |
| Total food intake (g) | 2199±16.04 | 2197±10.29 | 1809±7.77 | 1820±7.45 | >0.4967 | **<0.0001** | >0.3338 |

Note: Data are presented as mean ± SD. Two-way ANOVAs were used to assess the main effects of exercise and sex and interaction between exercise and sex on post-measures. \*Data for adrenal/body mass ratio are presented as $\times 10^{-4}$

### Exercised mice made fewer errors than sedentary controls during training

To determine if there were sex or exercise-training dependent differences in spatial learning during the training sessions (Figure 1) a repeated measures ANOVA was performed for the number of errors incurred during spatial training (Figure 1A). The analysis revealed an interaction between training day and exercise (*F* (4, 171) = 3.630, *p* = 0.0072). Exercised mice made significantly fewer errors than sedentary controls on training day one (*p* = 0.0026) and two (*p* = 0.0502). From the third day of training onwards, the exercised and sedentary groups had comparable errors before finding the escape hole. Regardless of exercise and training day, males made more errors than females (main effect of sex: *F* (1, 43) = 10.87, *p* = 0.002).

**Figure 1.**
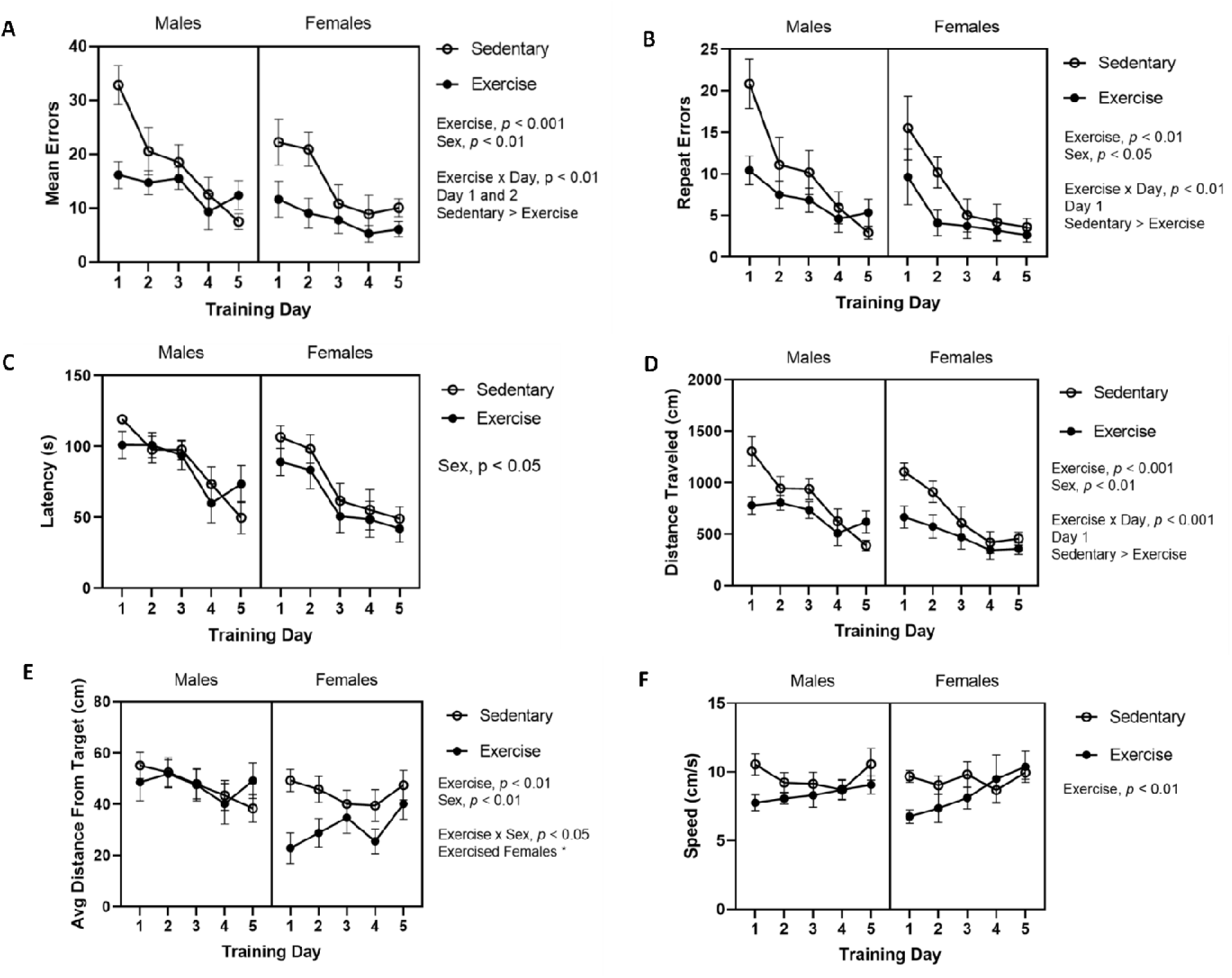
Effects of exercise on spatial learning in female and male mice in the Barnes maze. (A) The number of errors by training day, (B) The number of repeat errors by training day, (C) Latency by training day, (D) Distance travelled by training day, (E) Average distance from target by training day, and (F) Average speed by training day. Significant main (exercise, sex) and interaction (exercise by sex and exercise by day) effects are indicated on each figure. Error bars represent SEM. N = 12, 12, and 12, 11 mice for sedentary/exercised males and sedentary/exercised females, respectively.

We next examined differences in the number of repeated errors across training day (i.e. the number of times each incorrect target was visited more than once; Figure 1B), and we found a significant interaction between training day and exercise (F (4, 165) = 3.216, p < 0.0001), as well as a main effect of sex (F (1, 43) = 4.261, p = 0.0451), exercise (F (1, 43) = 8.181, *p* = 0.0065), and training day (*F* (2.933, 121.0) = 17.87, *p* < 0.0001). Exercised mice made fewer repeat errors than sedentary mice on the first day of training (*p* = 0.0252) and males made greater repeat errors than females (*p* = 0.0138).

A repeated measures ANOVA was performed on latency across training day (Figure 1C), which revealed a main effect of sex, with males having a longer latency than females (*F*(1, 43) = 10.97, *p* = 0.0019). There was also a significant effect of training day (*F* (3.353, 144.2) = 18.43, *p* < 0.001). A significant difference in latency was observed between training day one and three (*p* = 0.0004), which sustained significance from day one to day four (p < 0.0001), as well as day one to day five (*p* < 0.0001). There was no significant difference in latency between day one and day two (*p* = 0.6122).

### Exercised females spent more time near the escape hole

A repeated measures ANOVA was performed on the distance of path travelled across training days (Figure 1D). The analysis showed an interaction between training day and exercise (*F* (4, 165) = 3.551, *p* = 0.0083), as well as a main effect of training day (*F* (3.396, 140.1) = 16.96, *p* < 0.0001), sex (*F* (1, 43) = 12.42, *p* = 0.001), and exercise (*F* (1, 43) = 14.26, *p* = 0.0005). Exercised mice travelled a shorter distance compared to sedentary controls on training day one (*p* = 0.0006) and males travelled a farther distance than females before finding the escape hole.

A repeated measures ANOVA was performed on the average distance from the escape hole across training days (Figure 1E), which showed an interaction between sex and exercise (*F* (1, 43) = 5.897, *p* = 0.0194), as well as a main effect of sex (*F* (1, 43) = 11.54, *p* = 0.0015), and exercise (*F* (1, 43) = 5.845, *p* = 0.0199). The analysis revealed no effect of training day. Collapsing across all training days, exercised females traveled a shorter distance to the escape hole than exercised males (p < 0.0001), sedentary females (0.0184), and sedentary males (p < 0.0001).

To determine whether latencies or errors were affected by systematic differences in the speeds of the groups (Figure 1F), we analysed the average speed for each group on each training day. The analysis demonstrated that exercised mice moved slower than their sedentary counterparts, regardless of sex (main effect of exercise: *F* (1, 43) = 7.360, *p* = 0.0096).

### Exercised mice demonstrated greater use of spatial strategy

We examined whether there were differences in navigational strategy between the groups (Figure 2A-D) by determining the proportion of each strategy used across all five training days. We found that males used a random strategy significantly more often than females (main effect of sex: *F* (1, 43) = 11.32, *p* = 0.0016; Figure 2E) while females used a spatial strategy more often than males (effect of sex (*F* (1, 43) = 14.33, *p* =0.0005; Figure 2F). The proportion of animals using a spatial strategy was affected by exercise (*F* (1, 43) = 8.463, *p* =0.0057) with exercised mice using a spatial strategy more often than sedentary mice. While this effect seemed to be driven by female exercised mice, the interaction between exercise and sex was not significant (*F* (1, 43) = 1.729, *p* = 0.1955).

**Figure 2.**
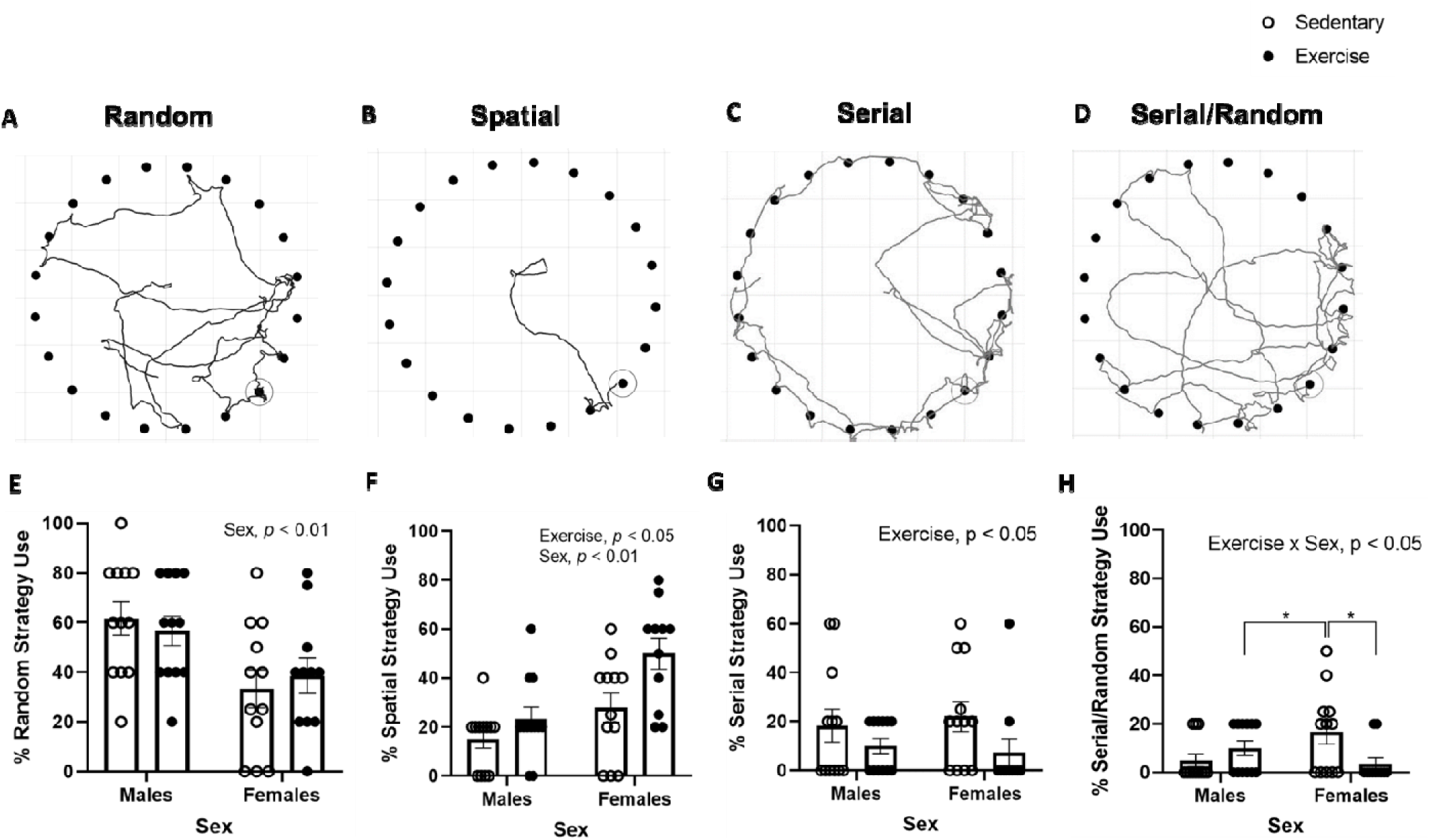
Effect of exercise training on strategy use in the Barnes maze in female and male mice. Example diagrams illustrating a (A) random, (B) spatial, (C) serial, and (D) combination serial/random search strategy. Percentage of sedentary and exercised mice employing a (E) random, (F) spatial, (G) serial, and (H) serial/random strategies during the learning phase in the Barnes maze. Significant main (exercise, sex) and interaction (exercise by sex) effects are indicated on each figure. Lines indicate significant differences between groups when the interaction (treatment x sex) was significant. Error bars represent SEM. N = 12, 12, and 12, 11 mice for sedentary/exercised males and sedentary/exercised females, respectively

However, exercised females did indeed accumulate a shorter distance to the escape hole compared to all other groups, thereby indicating a potential benefit from the exercise intervention that was modulated by sex. Exercised mice also used a serial strategy less often than sedentary controls (*F* (1, 43) = 4.343, *p* = 0.0431; Figure 2G). Finally, when comparing serial/random strategy use (i.e. the combination of both strategies employed during the trial), we found an interaction between exercise and sex (*F* (1, 43) = 6.867, *p* = 0.0121; Figure 2H) with sedentary females using this combination strategy significantly more than both sedentary males (*p* = 0.0392) and exercised females (*p* = 0.0225).

### Exercised mice made fewer errors during the probe trial

Following training, mice were tested in a probe trial to examine latency to correct hole, correct target visits, errors, and time spent in the target quadrant (Figure 3). There were no significant differences in latency or correct target visits during the probe. However, there was a significant effect of exercise on the number of errors (Figure 3A), with exercised mice visiting fewer incorrect targets during the probe session compared to sedentary mice (main effect of exercise: *F* (1, 43) = 5.187, *p* = 0.0278). There was a significant effect of sex on the time spent in the target quadrant, with females spending more time in this quadrant than males (*F* (1, 43) = 5.502, *p* = 0.0237; Figure 3B). Similarly, we also found that females spent significantly more time near the escape hole (*F* (1, 43) = 9.149, *p* = 0.0042; Figure 3C).

**Figure 3.**
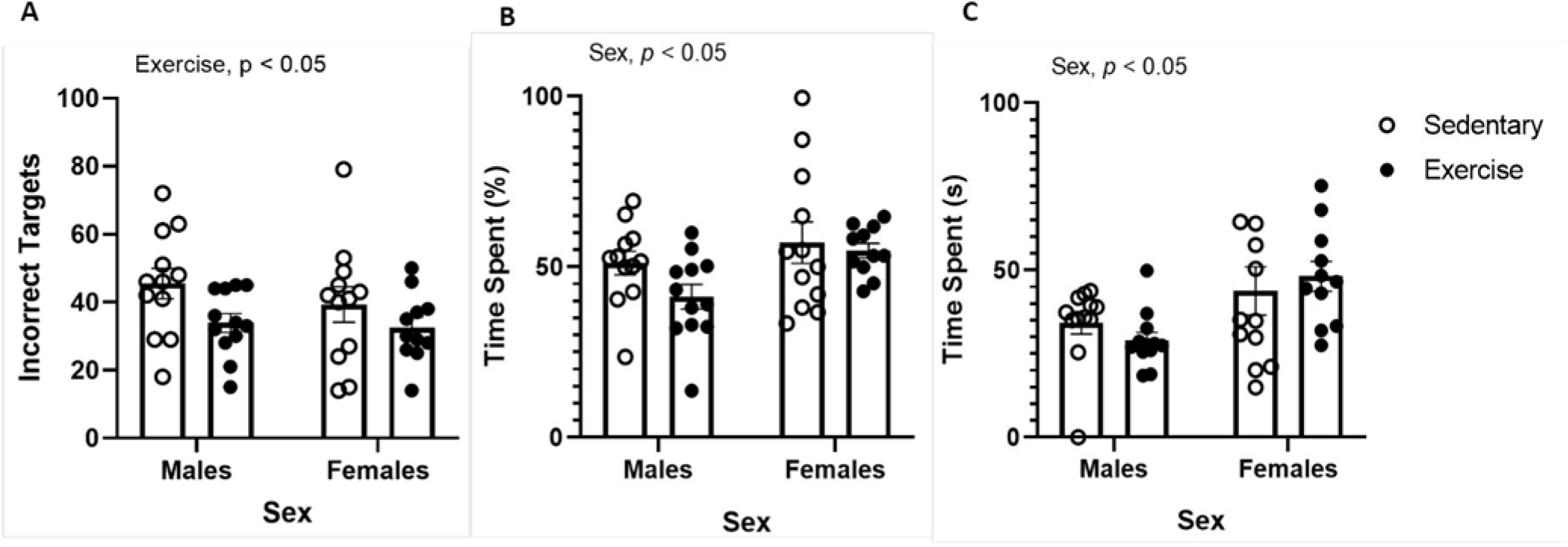
Effects of exercise on spatial memory performance in female and male mice during the probe session of the Barnes maze. (A) number of incorrect targets visited, effect of exercise (p < 0.05); (B) percent time spent in target quadrant, effect of sex (p < 0.05); (C) time spent near the escape hole, effect of sex (p < 0.05). Significant main (exercise, sex) effects are indicated on each figure. Error bars represent SEM. N = 12, 12, and 12, 11 mice for sedentary/exercised males and sedentary/exercised females, respectively

### Exercised mice had less WFA expression in the dorsal dentate gyrus compared to sedentary controls

To determine whether there were exercise or sex-dependent effects on the number of perineuronal nets, we examined WFA expression in the dorsal and ventral dentate gyrus (Figure 4A-C). We found that exercised mice had less WFA expression compared to sedentary mice in the dorsal region of the dentate gyrus (main effect of exercise: *F* (1, 44) = 4.236 *p* = 0.0455; Figure 4B). There were no significant effects of exercise, sex, nor an interaction in WFA expression in the ventral dentate gyrus (all *p*’s > 0.05; Figure 4C).

**Figure 4.**
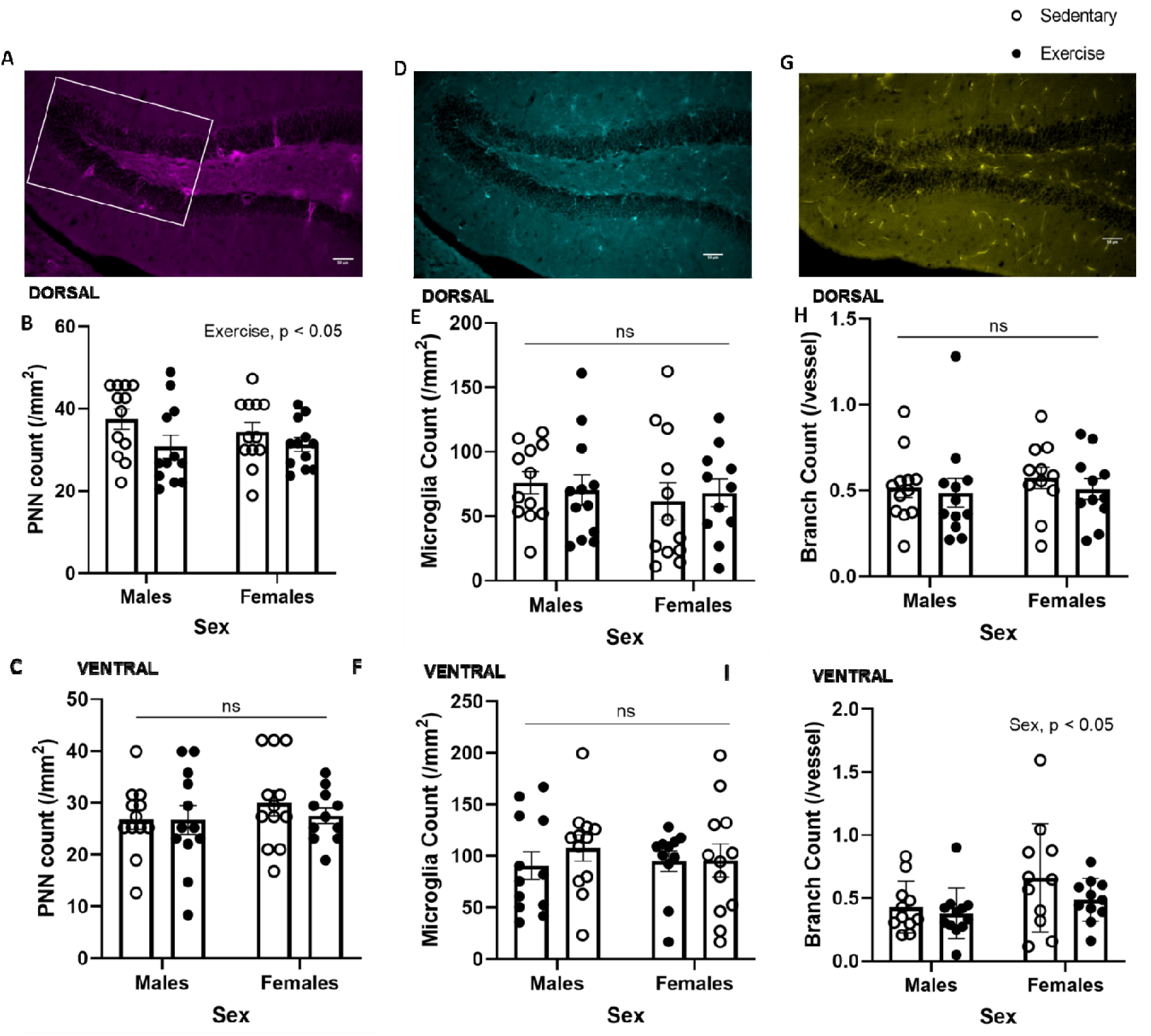
Effect of exercise training on PNN, microglia, and vascular branch counts in female and male mice. (A) PNN count in the (B) dorsal and (C) ventral dentate gyrus. (D) Microglia count in the (E) dorsal and (F) ventral dentate gyrus. (G) Vascular branch count in the (H) dorsal and (I) ventral dentate gyrus. Significant main (exercise, sex) effects are indicated on each figure (ns, indicated not significant). Error bars represent SEM. White box represents approximate size and placement of the region of interest. N = 12, 12, and 12, 11 mice for sedentary/exercised males and sedentary/exercised females, respectively. Four dorsal and three ventral images were used for each mouse brain.

### CD-31 and Iba-1 quantity were not affected by exercise

We determined the effects of exercise on microglia (using Iba-1) and blood vessels (using CD-31) in the dorsal and ventral dentate gyrus (Figure 4D-I). There were no significant effects of sex or exercise, nor any interactions (all p’s > 0.05) on Iba-1 counts in the dorsal or ventral dentate gyrus (Figure 4E-F). We also examined changes in blood vessel counts and vessel morphology (i.e., number of vessels, number of branches per vessel, and longest continuous section of each vessel) using CD31. There was a significant effect of sex on branch counts in the ventral dentate gyrus, with females having more branching per vessel than males (*F* (1, 41) = 4.426, *p* = 0.0416; Figure 4I), which was not significant in the dorsal region (Figure 4H). There were no significant effects of exercise, sex, nor any interactions regarding vessel lengths or number of vessels in either region (all *p*’s > 0.05).

We also investigated whether exercise training resulted in changes in microglia morphology (Figure 5A-C). There were no differences in morphological proportions in the dorsal dentate gyrus between any of the groups (all p’s > 0.05; Figure 5D). In the ventral dentate gyrus, there was a significant interaction between morphology type and exercise (*F* (2, 90) = 4.982, *p* = 0.0089). In sedentary mice, there were more thin and stout microglia compared to ameboid microglia in the ventral region. In the exercised animals, the proportions were shifted with similar levels of thin and ameboid microglia (Figure 5E).

**Figure 5.**
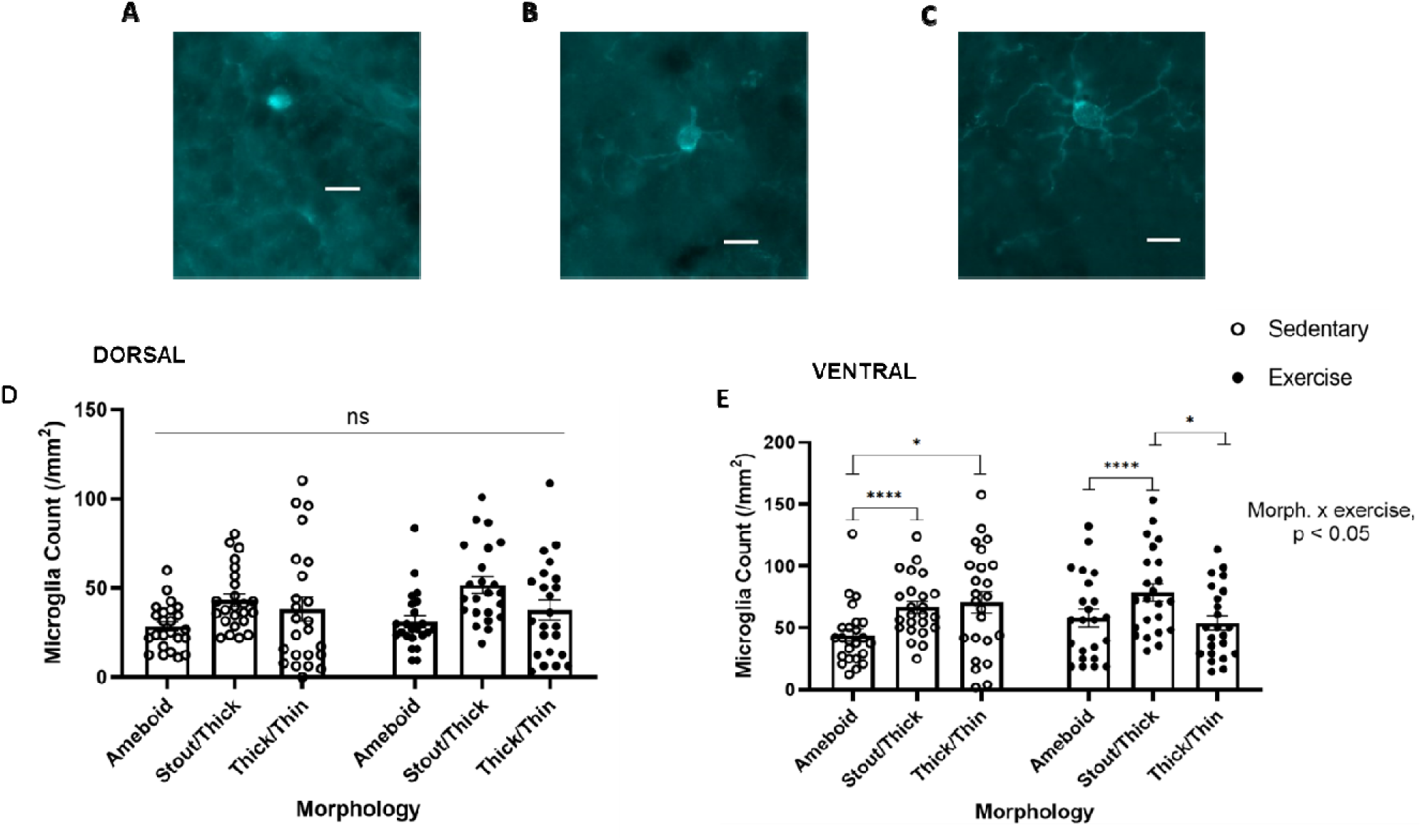
Representative images of different microglia morphology and effect of exercise training on microglia morphology in the dentate gyrus. (A) ameboid, (B) stout/thick, and (D) ramified/thin microglia. Scale bar represents 10 µm. Effect of exercise training on microglia morphology in the dorsal (D) and ventral dentate gyrus (E). No significant main (exercise, sex) or interaction (exercise by sex) effects were observed. The interaction between morphology and exercise was significant in the ventral region. Error bars represent SEM. N = 12, 12, and 12, 11 mice for sedentary/exercised males and sedentary/exercised females, respectively.

### Correlations between neurobiological measures and errors in the Barnes maze

We performed Pearson r correlations to determine whether there was a relationship between neurobiological and behavioural measures. There was a small, positive correlation between total errors during training and dorsal WFA expression amongst the whole group (*r* = 0.2876, *p* = 0.0500; Figure 6A), but no significant relationship existed within any subgroups (sex or exercise).

**Figure 6.**
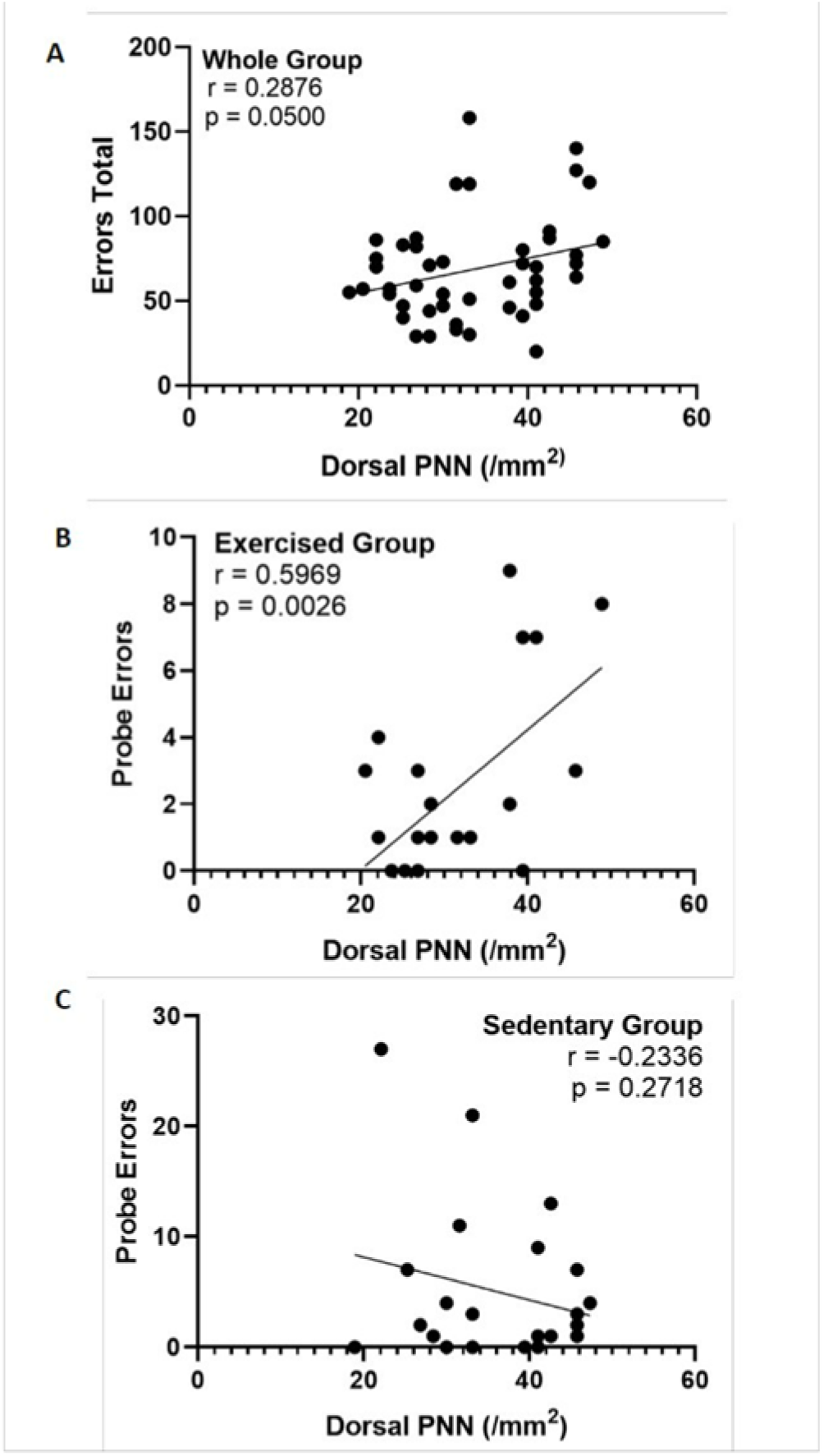
Correlations between PNN expression and errors during learning and probe. (A) Total errors during learning were correlated with the number of dorsal (r = 0.2876, p = 0.05); (B) dorsal PNNs and probe errors were correlated among the exercised group (r = 0.5969, p = 0.0026) but not among the sedentary group (C) (r = -0.2336, p > 0.05). N = 12, 12, and 12, 11 mice for sedentary/exercised males and sedentary/exercised females, respectively.

We found a significant positive correlation between dorsal WFA expression and errors before finding the escape hole during the probe among the exercised group (*r* = 0.5969, *p* = 0.0026, Figure 6B), with the correlation similar among the exercised females (*r* = 0.6451, *p* = 0.0321) and exercised males (*r* = 0.6264, *p* = 0.0293) but not in the sedentary group (*r* = -0.2336, *p* = 0.2718; Figure 6C). There were no significant correlations between dorsal WFA and correct target visits, or time spent in the target quadrant during the probe, amongst the whole group or otherwise.

In the ventral region, there were no significant correlations between WFA and total errors during training. However, there was a moderate negative correlation between ventral WFA expression and average repeat targets (data not shown) among the sedentary group (*r* = -0.4254, *p* = 0.0392), which was strongly driven by the sedentary males (*r* = -0.6858, *p* = 0.0146) as the sedentary females demonstrated no relationship (*r* = -0.1196, *p* = 0.7112) nor did either exercised group (exercised females: *r* = -0.2357, *p* = 0.4854; exercised males: *r* = 0.2652, *p* = 0.4049).

## Discussion

The purpose of this study was to determine the sex-specific effects of exercise training on spatial memory and neurobiological outcomes, as measured by microglia, PNNs, and vascularisation in the dentate gyrus of the hippocampus. Our findings indicate that in healthy adult mice, exercise improves spatial learning in both sexes, and this is associated with a reduction in PNNs but not with changes in vascularisation or microglia numbers within the dentate gyrus. We found that female mice performed better on the Barnes maze during training and probe trials, and we found sex differences in strategy use. Our results however indicate that treadmill exercise benefits spatial memory in both sexes and this is associated with changes in the extracellular matrix in the hippocampus.

### Exercise improved spatial learning and memory in both sexes

There has been ample previous research relating the benefits of exercise to improved learning and memory (Li et al., 2013; Xiong et al., 2015; Hwang et al., 2016). However, most of the past research has focused on males. We found that treadmill exercising adult mice, regardless of sex showed improved spatial learning and memory assessed with the Barnes maze. Previous work in adult male (van Praag et al., 2005) and female (van Praag et al., 1999) C57BL/6 mice using voluntary wheel running also showed improvements in spatial memory, tested with the Morris water maze. Our use of the Barnes maze, as opposed to the Morris water maze is an overall strength in the context of our study, as the swimming component of the Morris water maze may bias results towards animals with higher fitness. Our results demonstrate that the exercise-induced benefit was most prevalent during the first two training days, during which time the exercised mice had significantly fewer errors, thereby indicating hastened spatial learning compared to sedentary controls. This may suggest that the exercised mice created a more stable representation of the novel maze environment from the initial habituation session. This is also supported by the finding that the exercise trained mice made fewer repeat errors on the first day of training, which may also indicate enhanced working memory. We also found that exercised mice travelled less during the first day of training and had shorter total distances to the target hole across training days (Figure 2 and 3). These findings suggest that exercise might not only improve memory, but also speed of learning (van Praag et al., 1999; Winter et al., 2007; Ji et al., 2015; Jacotte-Simancas et al., 2013). In addition, in the current study, exercise also improved recall ability, as the exercised mice made fewer errors during the probe session compared to their sedentary counterparts. Lastly, the enhanced performance in the Barnes maze among the exercised group does not seem to be the result of enhanced locomotion or fitness, as the exercised group demonstrated slower locomotion during training.

### Females demonstrated enhanced spatial memory compared to males

Our results showed that females demonstrated improved spatial learning compared to their male counterparts, as evidenced by fewer total errors observed during training among the female mice. Although contrary to the generally accepted notion of male-favoured spatial memory performance that is heralded in human literature (Iachini et al. 2005; Lejbak et al., 2011; Yuan et al., 2019), and mirrored in animal research (Gresack & Frick, 2003; Safari et al., 2021), this study demonstrated female-favoured performance in spatial navigation, as measured by fewer errors and increased spatial strategy preference among female mice. Female preference for spatial strategy compared to males is consistent with research conducted in degus using the Barnes maze (Popović et al., 2010), and though not significant in this investigation, the preference for spatial strategy use appeared to be driven by the exercised females. Exercised females also had the shortest average distance to the escape hole during training. Together, these findings indicate that exercise may pose a greater cognitive benefit for female compared to male mice. Previous work has shown that the distal placement of spatial cues in relation to the navigator can affect performance and that this can differ by sex (Saucier et al., 2007). Specifically, females tend to benefit from peri-personal cues (spatial markers that are close to the navigator), while males tend to benefit more so from extra-personal cues (spatial markers that are more distant from the navigator; Saucier et al., 2007). As such, the placement of the spatial cues may have provided greater benefit to females compared to males, and this benefit, combined with that of exercise, may have had a cumulative effect to provide exercised females with the greatest advantage in this spatial memory task.

### Exercise reduced hippocampal perineuronal nets in both sexes

Our findings showed that exercise reduces the number of PNNs in the dorsal, but not the ventral dentate gyrus in female and male mice. Although body composition was not affected by our exercise regimen, previous work has found that the same exercise protocol increases BDNF content in male mice without affecting body mass (Baranowski et al., 2023). Some studies have reported reductions in body mass with exercise (Castellani et al., 2014) but body adaptations to exercise vary and depend on the age and health of the mouse model. In the current study, we observed improved spatial learning and memory in exercised mice, accompanied by changes in perineuronal nets, suggesting that alterations in body composition or mass may not be necessary for brain changes to occur. Previous studies have suggested that the dorsal hippocampus is more involved with spatial memory, whereas the ventral hippocampus is more involved with motivated behaviours and stress reactivity (Kheirbek & Hen, 2011). Our results are consistent with the notion of this distinct bifunctionality within the hippocampus, as perineuronal nets decreased only in the dorsal region. This exercise-induced PNN reduction is also consistent with previous work conducted in female rats using voluntary wheel running (Smith et al., 2015). Using a controlled exercise regimen, we showed that reduction in PNNs due to exercise was associated with enhanced spatial memory performance in both sexes, as total errors were correlated with the number of PNNs during training. Furthermore, greater PNNs was also associated with poorer spatial memory performance (indicated by errors) during the probe session among the exercised group.

To our knowledge, there has been no work regarding the effects of exercise on PNNs with respect to spatial memory outcomes, nor with respect to possible sex differences. However, a review by Bozzelli and colleagues (2018) proposes a hypothetical model of PNN expression with respect to synaptic plasticity and synaptic stability. The model suggests that synaptic stability increases with PNN expression, which also occurs as a function of age. Consequently, synaptic plasticity is slowly replaced by synaptic stability through PNN deposition as age increases (Bozzelli et al., 2018). Certain interventions, such as exercise, can decrease PNNs within an optimal range that has not yet been defined or quantified. Our results are consistent with this theoretical framework as we demonstrated lower PNNs within the dentate gyrus, which is indicative of increased synaptic plasticity, as generally suggested by this framework. Synaptic plasticity is conducive to learning and memory, as the latter processes require the formation of new neural networks (Stuchlik, 2014), which may be aided by reduced PNNs (Carstens et al., 2016). Correspondingly, enhanced synaptic plasticity, as indicated by exercise-induced PNN reductions in the dentate gyrus, is also consistent with our observation of improved learning among the exercised group during training. Further, correlation analyses were conducted to explore possible relationships between PNNs and memory performance. A significant positive relationship between PNN count and probe errors was only observed among the exercised group, which may indicate that the role of PNNs are affected by exercise. Previous work has found that PNN degradation is associated with improved object recognition memory in male mice (Romberg et al., 2013) but also impairments in object recognition memory in macaques (Gray et al., 2023). This suggests there is a complex relationship between PNN expression and memory when factors such as type of memory, brain region, as well as stage of learning and memory (acquisition, recall, retention, and reconsolidation) are considered (Gogolla et al., 2009; Romberg et al., 2013; Shi et al., 2019; Christensen et al., 2021). Finally, one last factor to be considered is the PNN marker being used. Wisteria floribunda agglutinin (WFA) is the most commonly used marker for PNNs (Härtig et al., 2022). However, recent work has demonstrated that WFA does not bind to all PNNs, nor do other common PNN markers such as those binding aggrecan or chondroitin sulphate proteoglycan link proteins, and this depends on brain region (Hartig et al., 2022). Further work is needed to better understand the different markers and the potential functional implication of different populations of PNNs.

### Exercise did not affect vascularisation and had limited effects on microglia

Microglia have many integral functions within the brain that relate to neuro-immunity, neuroinflammation, neuroplastic processes, and learning and memory (Cherry et al., 2014; Czerniawski et al., 2014). Ameboid and stout or thick microglia are typically associated with phagocytosis and increased neuroinflammation (although see Paolicelli et al., 2022), which can contribute to impaired cognition (Cherry et al., 2014; Czerniawski et al., 2014). Treadmill exercise did not affect the number of microglia in the dentate gyrus. However, we did find that the proportions of microglia type (ameboid, stout/thick, and thick/thin) were regulated by exercise. Exercised mice had a reduction in the proportion of thin microglia. Although we expected that exercise would reduce neuroinflammation, indicated by a decrease in microglia numbers or shifts in microglia morphology, our results do not support this hypothesis. A large portion of previous work has been conducted using models of neurological disease and has shown that exercise reduces hippocampal neuroinflammation in these contexts (Nichol et al., 2008; Ke et al., 2011; Tapia-Rojas et al., 2016; Zhang et al., 2018; Wang et al., 2023). Our work adds to the limited research conducted in healthy models and suggests that in healthy adult mice, chronic treadmill exercise does not measurably regulate microglia count in the dentate gyrus. Our results suggest exercise has a different mechanism of action in the brain of healthy individuals and that changes in microglia numbers and morphology may not be involved in the beneficial effects of exercise on learning and memory in a healthy model. However, further work using other measures of neuroinflammation are needed in healthy animal models. Microglia release both pro- and anti-inflammatory cytokines (Ferro et al., 2021) and exercise is both a pro and anti-inflammatory process (Docherty et al., 2022). A healthy brain demonstrates a coexistence and balance of pro- and anti-inflammatory cytokines (Crain et al., 2013). Consequently, it is possible that exercise regulates this balance without affecting microglia numbers or morphology.

Although there is evidence of exercise-induced angiogenesis in the body (Prior et al., 2004), there is limited work regarding this process in the brain. We used a direct approach through the use of CD31 and measurement of blood vessel length, count, and branching. Despite this, we did not find an exercise-induced increase in angiogenesis within the dentate gyrus in adult female and male mice. Previous work has demonstrated increased blood volume in this region in mice and humans (measured by magnetic resonance imaging), as well as an increase in vascular endothelial growth factor in mice using other types of exercise such as high intensity interval training (HIIT) and voluntary wheel running (sex was not reported; Pereira et al., 2007; Morland et al., 2017). However, these methods do not necessarily suggest vascular remodelling. In contrast, we examined structural vascular changes in relation to learning and memory. Altogether, our results along with previous studies suggest that exercise may remodel the brain through changes in vascular function but not necessarily structure.

### Limitations

A common debate in exercise research pertains to the benefits and drawbacks of treadmill exercise and voluntary wheel running. While treadmill running may be more stressful than voluntary wheel running, it is generally well tolerated, and protocols are in place to reduce stress. Prior to any treadmill training mice are habituated to the treadmill environment to reduce stress. As a marker of stress, we measured adrenal mass and found no differences between experimental groups. The advantage of using treadmill running is that it allows for a controlled intensity and duration of exercise thus eliminating any variability in these parameters with voluntary wheel activity. For example, previous work from our group has demonstrated that female mice may run as little as 0.33 km per night and as high as 1.2 km per night (Mohammad et al., 2022). Further, and relevant to our goal of comparing adaptations between sexes, research indicates that there are notable sex-specific differences in voluntary wheel running behaviour in mice. In C57BL/6J mice, females have been reported to run on average 40% further than males, at greater speeds, and for longer durations (De Bono et al., 2006; White et al., 2016). Accordingly, although treadmill running may be more stressful compared to voluntary wheel running, in the context of this study which compared sex differences in neural mechanisms and cognition, it was necessary to use this exercise regimen to control for running distance and intensity.

Here we used the Barnes maze to assess changes in spatial learning and memory because it provides a variety of behavioural data from which extensive knowledge can be gleaned, including different types of errors, latency, distance, speed, and strategy. In contrast with one-trial paradigms, a crucial strength of the Barnes maze test is the tracking of improvement in performance over several days, which allowed us to capture the process of learning rather than working or short-term memory. The ability to capture the process of learning comes at a cost to time as the entire process from the habituation phase to the final probe session takes one week to perform. This timing did not allow us to incorporate additional cognitive tests, but future work should assess other types of memory in response to exercise in both sexes.

Although we have demonstrated that exercise reduces PNNs in the dentate gyrus of adult mice and demonstrated a that a relationship exists between PNNs and spatial learning and memory, our work cannot determine whether the reduction in PNNs directly affected memory outcomes. Additionally, comparison of these findings with studies that have investigated cognitive effects of PNN degradation through chABC administration should be done so with caution, as chABC is a bacterial enzyme that is used to pharmacologically degrade PNNs to produce phenotypes consistent with juvenile-like plasticity (Hettiaratchi et al., 2019; Willis et al., 2022), beyond reductions that have been observed through exercise. Further work regarding the role of exercise-induced reductions in PNNs in relation to cognitive outcomes is required. Development of a method that blocks PNN degradation would be beneficial in observing the role of this process in learning and memory.

Finally, with respect to angiogenic outcomes, it was unexpected that we were unable to find measurable vascular changes in branch counts and branch lengths. The lack of changes we observed with respect to vascular density do not necessarily preclude the occurrence of angiogenesis, as other related phenomena may have prevented observable changes, such as vascular pruning or adaptational metabolic changes within the vascular tissue that were not otherwise accounted for.

## Conclusion

In conclusion, we found that endurance training reduces PNNs content in the dentate gyrus in both sexes and this may be related to enhanced spatial learning and memory performance. Surprisingly, microglia and vascular remodelling were not regulated by exercise, pointing to potential differences in the underlying mechanisms of action in the brain of exercise in healthy individuals, which highlights the novelty in our study. While we found that females performed better in the Barnes maze, this sex difference in spatial memory was not associated with changes in PNNs in the dentate gyrus. Further work is needed to elucidate underlying mechanisms of sex differences in spatial memory, including other neurobiological measures and brain regions.

## Acknowledgements

This research was supported by the Natural Sciences and Engineering Council of Canada to RM (RGPIN-2017-03904) and the Canada Research Chair Program to PDG. We thank the animal care staff at the Comparative Bioscience Facility at Brock University for technical assistance. AY is funded by an NSERC Alexander Graham Bell Canada Graduate Scholarship-Doctoral (CGS-D).

## Notes

### Competing Interest Statement

The authors have declared no competing interest.

### Summary of Updates

Discussion of the manuscript was updated. Figure 3 was revised.

